# Steady-state phosphorylation of SHIP1 by Lyn restricts macrophage activation in the absence of a phagocytic synapse

**DOI:** 10.1101/2025.04.16.648985

**Authors:** S. Erandika Senevirathne, William K. Kanagy, Richard T. Cattley, Whitney L. Swanson, Silvia Toledo Ramos, Myra G. Nunez, William F. Hawse, Tanya S. Freedman

## Abstract

Microscale engagement of the hemi-ITAM-containing receptor Dectin-1 by a fungus-derived particle initiates signaling through the Src-family kinases (SFKs) and Syk that leads to downstream Erk and Akt activation and the macrophage anti-microbial response. To minimize local tissue damage in the absence of a true pathogenic threat, macrophages must remain unresponsive to low-valency receptor ligation by soluble ligands from food or remote infection sites. To investigate how SFKs regulate macrophage sensitivity, we compared signaling in murine bone-marrow-derived macrophages (BMDMs) exposed to depleted zymosan, a particulate and highly multivalent Dectin-1 ligand, with signaling after pharmacologically induced SFK activation without receptor engagement or formation of a phagocytic synapse. We show that high-valency Dectin-1 engagement and formation of a phagocytic synapse restrict phosphorylation of the ITIM-associated phosphatase SHIP1 and promote Erk and Akt signaling. In contrast, SFK activation in the absence of a phagocytic synapse induces phosphorylation of SHIP1 and leads to dampened activation of Erk and Akt. Whereas multiple SFKs are capable of phosphorylating SHIP1 in principle, the SFK Lyn functions uniquely in maintaining steady-state phosphorylation of SHIP1 and setting tonic and induced levels of Erk and Akt phosphorylation. Consequently, Lyn has a special role in suppressing the Akt pathway and maintaining Erk/Akt pathway balance. Interestingly, formation of a phagocytic synapse circumvents this requirement for Lyn to maintain Erk/Akt balance. These findings highlight the unique function of Lyn in maintaining macrophage steady-state signaling and limiting pro-inflammatory responses to disorganized pathway activation and soluble microbial components.

## Introduction

Engagement of macrophage ITAM- and hemi-ITAM-coupled receptors (e.g., FcγR, Dectin-1) by highly multivalent ligands, such as antibody-opsonized pathogens or fungal-wall β-glucans, activates Src-family kinases (SFKs), which phosphorylate ITAMs and recruit the kinase Syk [1]. When Syk is bound to phosphorylated ITAM tyrosine (Y) residues, SFKs phosphorylate Y352 and Y348 its SH2-kinase linker to promote full Syk activation [2, 3]. Together, activated SFKs and Syk phosphorylate downstream effectors, including the second-messenger-generating enzymes phospholipase (PL)Cγ2 and phosphoinositide 3-kinase (PI3K). PLCγ2 hydrolyzes phosphatidylinositol 4,5-bisphosphate (PI(4,5)P_2_) to membrane-associated diacylglycerol and soluble inositol trisphosphate (IP_3_) to mobilize calcium and activate signaling through the small G-protein Ras and ultimately the kinases Erk1/2 [4]. PI3K phosphorylates PI(4,5)P_2_ to phosphatidylinositol 3,4,5-trisphosphate (PIP_3_), which recruits and allosterically activates PH-domain-containing kinases such as PDK1, mTORC2, and Akt [5, 6]. Phosphorylation of Akt by PDK1 and mTORC2 on threonine (T) 308 and serine (S) 473, respectively, activates Akt [7, 8]. PIP_3_ also mediates the recruitment and activation of PH-domain-containing proteins upstream of Erk1/2, including BTK and PLCγ2 [9, 10]. As antimicrobial byproducts can be toxic to neighboring tissues and drive autoimmune disease, activation of macrophages must be tightly regulated to minimize collateral damage [11-13]. To this end, SFKs also mediate immune suppression by phosphorylating ITIMs, which recruit cytosolic inositol phosphatases (e.g., SHIP1) and tyrosine phosphatases (e.g., SHP-1) to dampen Akt and Erk signaling by dephosphorylating PIP_3_ and tyrosine-phosphorylated proteins [14, 15].

Different types of myeloid cells and lymphocytes have distinct SFK expression profiles, and the contributions of different Src family members to cell function are not clear [16-18]. The SFK Lyn, which is expressed in myeloid and B cells, can play both positive and negative roles in immune regulation. Along with other SFKs, Lyn participates in pro-inflammatory ITAM signaling and the antimicrobial response, but a general negative-regulatory role is evidenced by the progression of chronic inflammation and autoimmunity in mice lacking Lyn expression [19, 20]. Lyn knockout (Lyn^KO^) mice have increased numbers of autoreactive B cells and pro-inflammatory myeloid cells and increased production of inflammatory cytokines and autoantibodies [21-23]; isolated Lyn^KO^ B cells have increased calcium mobilization [19, 20] and hyperresponsive Erk signaling [11, 22]. Crosslinking B-cell receptor (BCR) in Lyn^KO^ B cells leads to more Akt^S473^ phosphorylation than in wild-type (WT) cells [11]. These signaling abnormalities stem from impaired phosphorylation of ITIMs (CD22,

FcγRIIb) and ITIM-associated phosphatases (SHIP1, SHP-1) at steady state and in response to antigen encounter [20, 24-26]. In mast cells the function of Lyn appears to vary according to ITAM cluster size: low-valency FcεRI clustering promotes a pro-inflammatory function, whereas high-valency clustering promotes a suppressive function [27]. In addition to Lyn, macrophages express substantial amounts of the Src family members Hck, Fgr, and Fyn [17]. Previous studies show some functional redundancy between Lyn, Hck, and Fgr in FcγR-mediated phagocytosis and pro-inflammatory signaling [28, 29], but the degree of functional overlap within the Src family is not well understood.

In all cells SFK activity is tightly regulated by the opposing roles of the tyrosine kinase Csk and the membrane phosphatases CD45 and CD148. At steady state Csk phosphorylates Y507 in the SFK C-terminal inhibitory tail that nucleates an intra-molecular interaction with the SH2 domain to maintain an autoinhibited assembly [30]. In continuous competition with Csk, CD45 and CD148 dephosphorylate the SFK inhibitory tail, releasing autoinhibition and enabling activation-loop autophosphorylation [30]. Since CD45 and CD148 can also dephosphorylate the activation loop of SFKs, their steric exclusion from the phagocytic synapse promotes long-lived SFK activation and signal initiation at points of pathogenic contact [1, 31-33].

To investigate the roles of Lyn and the phagocytic synapse in macrophage signal initiation and negative feedback, we compared signaling triggered by a particulate (microscale) engagement of Dectin-1 with signaling triggered by pharmacological activation of SFKs in the absence of a phagocytic synapse. To activate the SFKs without formation of a phagocytic synapse, we used an analog of the kinase-inhibitor PP1 (3-IB-PP1) to specifically inhibit an analog-sensitive variant of Csk (Csk^AS^) [34-36]. Inhibiting Csk^AS^ blocks the dynamic equilibrium between Csk and CD45/CD148, leading to rapid SFK activation in the absence of a phagocytic synapse [34]. We have previously shown that receptor-independent activation of SFKs and downstream signaling can be achieved by pharmacologically inhibiting Csk, even in the absence of steric exclusion of CD45/148 [34].

Our studies reveal a unique role for Lyn in maintaining steady-state inhibitory signaling via SHIP1 phosphorylation, which limits accumulation of PIP_3_ and restricts Erk and Akt activation in macrophages. During SFK activation in the absence of a phagocytic synapse, Lyn is uniquely required to maintain Erk/Akt pathway balance by promoting Erk activation and inhibiting Akt activation. We demonstrate that disrupted basal inhibitory signaling and phospholipid homeostasis underlie this pathway dysregulation. Although other activated SFKs are intrinsically capable of phosphorylating SHIP1, this function fails to suppress Akt activation in the absence of a phagocytic synapse. These findings underscore the crucial function of Lyn in preventing spurious macrophage activation by maintaining basal signal homeostasis. Furthermore, our studies reveal that the phagocytic synapse obviates the requirement for Lyn to balance Erk and Akt signaling. Our data also suggest that SHIP1 is excluded from the phagocytic synapse, enabling signal amplification. These findings suggest that Lyn is a gate-keeper, preventing macrophage activation in response to nonparticulate stimuli or inappropriate SFK activation by maintaining a check on signaling in the absence of a direct encounter with a pathogenic cell.

## Results

### SHIP1 and SHP-1 restrain signaling downstream of disinhibited SFKs

To probe the role of the phagocytic synapse in SFK-initiated signaling, we first assessed signaling complexes initiated by SFKs in the presence and absence of receptor ligation. We performed imaging studies to quantify spatial differences in signaling initiated pharmacologically via the Csk^AS^ inhibitor 3-IB-PP1 compared to complexes formed around a particle of depleted zymosan, a yeast-derived β-glucan that signals specifically through Dectin-1. We visualized bone-marrow-derived macrophages (BMDMs) using total Lyn staining and used a direct substrate of the SFKs, phospho-Syk Y352 (pSyk^Y352^), as a reporter of signal activation. Unlike untreated controls **(Fig.1A)**, cells treated with 3-IB-PP1 developed puncta of phosphorylated Syk, dispersed across the cell **(Fig.1B)**. BMDMs treated with depleted zymosan, in contrast, signaled primarily at the phagocytic synapse **(Fig.1C)**. Although the larger puncta that form after 3-IB-PP1-treatment are most striking in the representative image, most of the puncta were on the nanoscale (between 0.1 and 1 µm^2^) and distributed throughout the cell **(Fig.1D)**. As observed previously [31], treatment with depleted zymosan induced highly localized phosphorylation of Syk^Y352^, arranged semi-contiguously around a microscale (∼70 µm^2^) phagosome. This comparison of organized/microscale vs. distributed/nanoscale signaling complexes allows us to assess quantitative and qualitative differences in signaling in the presence or absence of a phagocytic synapse.

**Fig.1:**
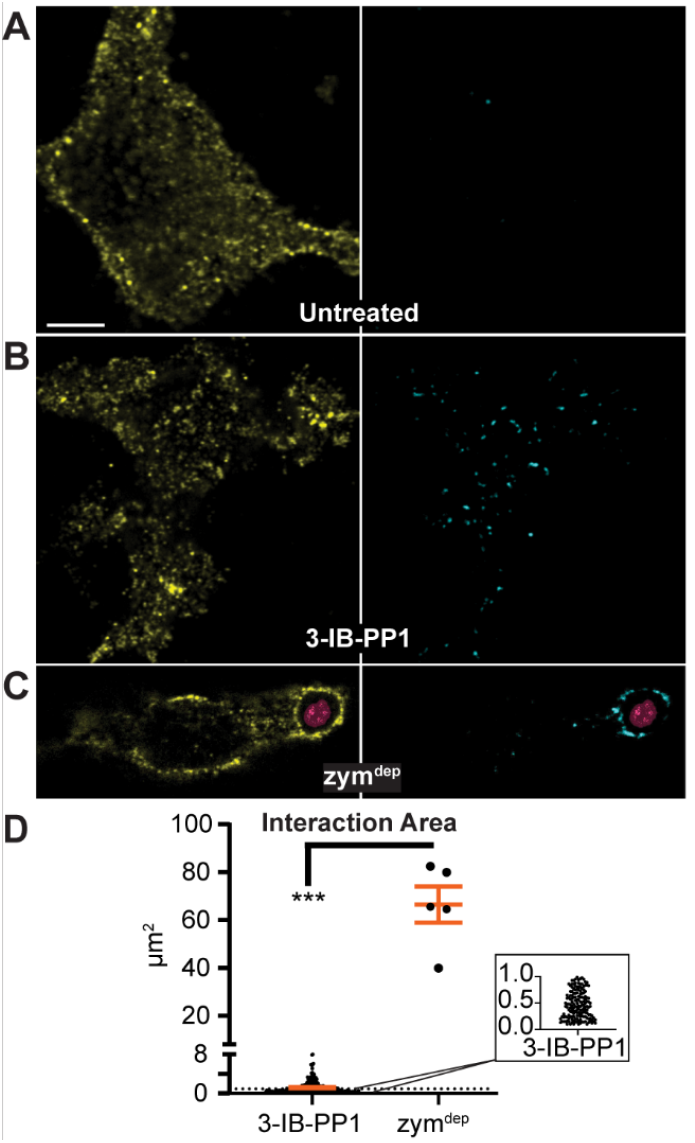
Pharmacological SFK activation induces only dispersed nanoclustering of signaling complexes. **(A-C)** Representative confocal micrographs showing a single z slice (∼280 nm) of IFN-γ-primed Csk^AS^ BMDMs stained for total Lyn (yellow) and pSyk^Y352^ (cyan) after 3 min treatments with **(A)** medium only (untreated), **(B)** 3-IB-PP1, or **(C)** depleted zy-mosan (zym^dep^). Fluorescence in the DAPI channel shows an internalized zym^dep^ (magenta). Scale bars: 5 μm **(D)** Interaction area for pSyk^Y352^ clusters formed spontaneously during 3-IB-PP1 treatment and semi-contiguous interaction area per phagocytosed zym^dep^ particle. Points are single-punctum values combined from 5 cells, shown with standard error of the mean (SEM). Significance (Sig.) in all panels assessed via nonparametric t test and Kolmogorov-Smirnov test: ^***^P = 0.0001. Inset: view of the lower range of 3-IB-PP1-induced pSyk clustering, showing the preponderance of nanoclusters within the distribution.

Consistent with our previous, qualitative studies [34], pharmacological SFK activation with 3-IB-PP1 produced more (35x) interdomain-B-phosphorylated Syk than did Dectin-1 ligation by depleted zymosan **(Fig.2A)**. Similarly, phosphorylation of PLCγ2 **(Fig.2B)** and PI3K **(Fig.2C)** was strongly induced by pharmacological SFK activation compared to treatment with depleted zymosan. Despite IFN-γ priming, which upregulates the Src-family kinase Lyn [34], among other pro-inflammatory factors [37], 3-IB-PP1-treated BMDMs failed to induce robust Ras nucleotide exchange **(Fig.2D)**, resulting in weaker Erk phosphorylation than cells treated with depleted zymosan **(Fig.2E)**. Akt phosphorylation was also impaired in the absence of particulate receptor engagement **(Fig.2F)**.

**Fig.2:**
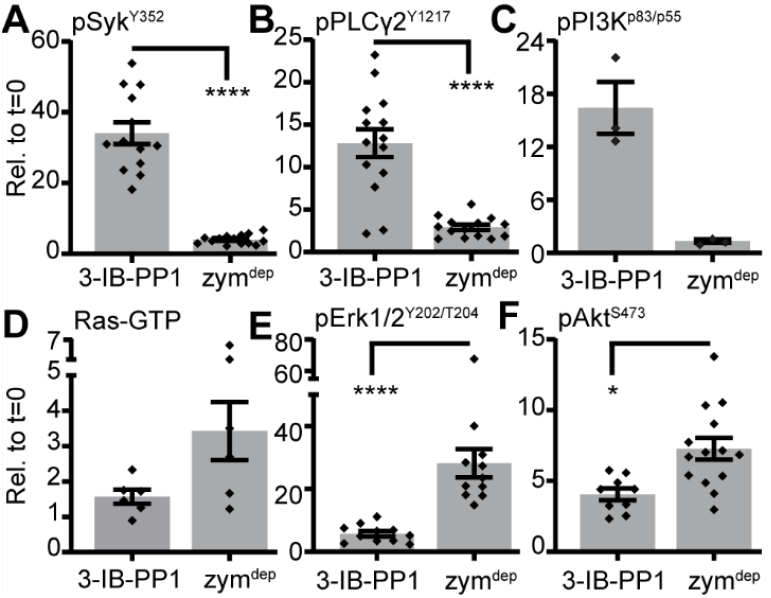
Signaling is amplified more effectively via particle engagement than via SFK activation alone. Densitometry quantification of protein species in immunoblotted lysates from IFN-γ-primed Csk^AS^ BMDMs after 5 min treatment with 3-IB-PP1 or zym^dep^ probed by immunoblot for **(A)** pSyk^Y352^, **(B)** pPLCγ2^Y1217^, **(C)** pPI3K^p85/p55^ **(D)** immunoprecipitated Ras-GTP, **(E)** pErk1/2^Y202/T204^, and **(F)** pAkt^S473^. Error: SEM; n=9-14 for 3-IB-PP1 and 11-15 for zym^dep^. Sig. assessed via non-parametric t test and Kolmogorov-Smirnov test: ^*^P = 0.0216, ^***^P = 0.0007, ^**^P < 0.0001.

Despite the weak upstream signaling induced by microscale Dectin-1 ligation, Ras-GTP, Erk, and Akt activation were robustly amplified. This disparity in signal amplification between 3-IB-PP1 and depleted zymosan also included differential phosphorylation of the inhibitory ITIM-associated inositol phosphatase SHIP1 and tyrosine phosphatase SHP-1. Although treatment with 3-IB-PP1 induced strong phosphorylation of Syk and PLCγ2, it also induced a 7-fold increase in SHIP1 phosphorylation **(Fig.3A)** and a 10-fold increase in SHP-1 phosphorylation **(Fig.3B)**. In contrast, the phagocytic synapse around particles of depleted zymosan protected SHIP1 and SHP-1 from phosphorylation above steady-state levels, indicating that these phosphatases could not access the signalosome at the membrane.

**Fig.3:**
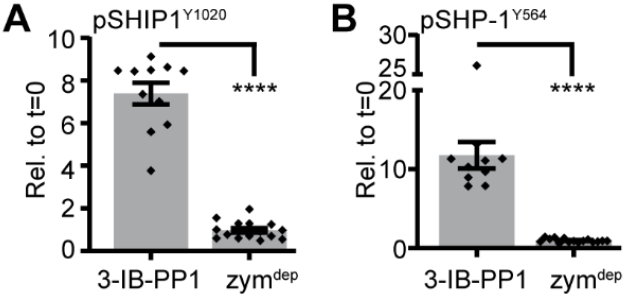
The phagocytic synapse restricts induced phosphorylation of inositol and protein phosphatases. Densitometry quantification of protein species in lysates from IFN-γ-primed Csk^AS^ BMDMs after 5 min treatment with 3-IB-PP1 or zym^dep^ probed by immunoblot for **(A)** pSHIP1^Y1020^ **(B)** pSHP-1^Y564^. Error: SEM; n=9-14 for 3-IB-PP1 and 11-15 for zym^dep^. Sig. assessed via nonparametric t test and Kolmogorov-Smirnov test: ^****^P < 0.0001.

The activation of SFKs in the absence of a phagocytic synapse forms dispersed nanoscale signaling puncta, mirroring a distribution of receptor nanoclustering by soluble ligands. Despite strong membrane-proximal signaling in the absence of a phagocytic synapse, a signaling bottleneck occurs between PLCγ2 and PI3K, as evidenced by impaired generation of Ras-GTP, pErk, and pAkt.

### Lyn maintains steady-state homeostasis by promoting SHIP1 phosphorylation and restricting enhanced Erk and Akt phosphorylation

Low levels of steady-state SFK activity in myeloid cells tune their sensitivity and enable rapid responses through ITAM, TLR, and cytokine receptor pathways [16, 38, 39]. We have previously shown that inflammation-dependent upregulation of Lyn sensitizes macrophages to receptor-independent signal propagation [34]. Since Lyn contributes to both positive and negative signaling downstream of ITAMs [27, 40, 41], we first examined the contributions of Lyn to steady-state macrophage signaling. We assessed protein phosphorylation in BMDMs primed overnight with a low dose of IFN-γ to upregulate Lyn expression and maximize the role of Lyn in maintaining the regulation of this sensitized steady state [34].

Steady-state phosphorylation of Syk^Y352^ was 3x lower in Lyn^KO^ BMDMs than in Lyn-expressing (Lyn^+/+^) BMDMs **(Fig.4A)**. Despite this potential loss of initiating signaling, phosphorylation of downstream mediators was elevated in Lyn^KO^ BMDMs, with 1.5x higher pErk1/2^T202/Y204^ **(Fig.4B)** and 3x higher pAkt^S473^ **(Fig.4C)**.

**Fig.4:**
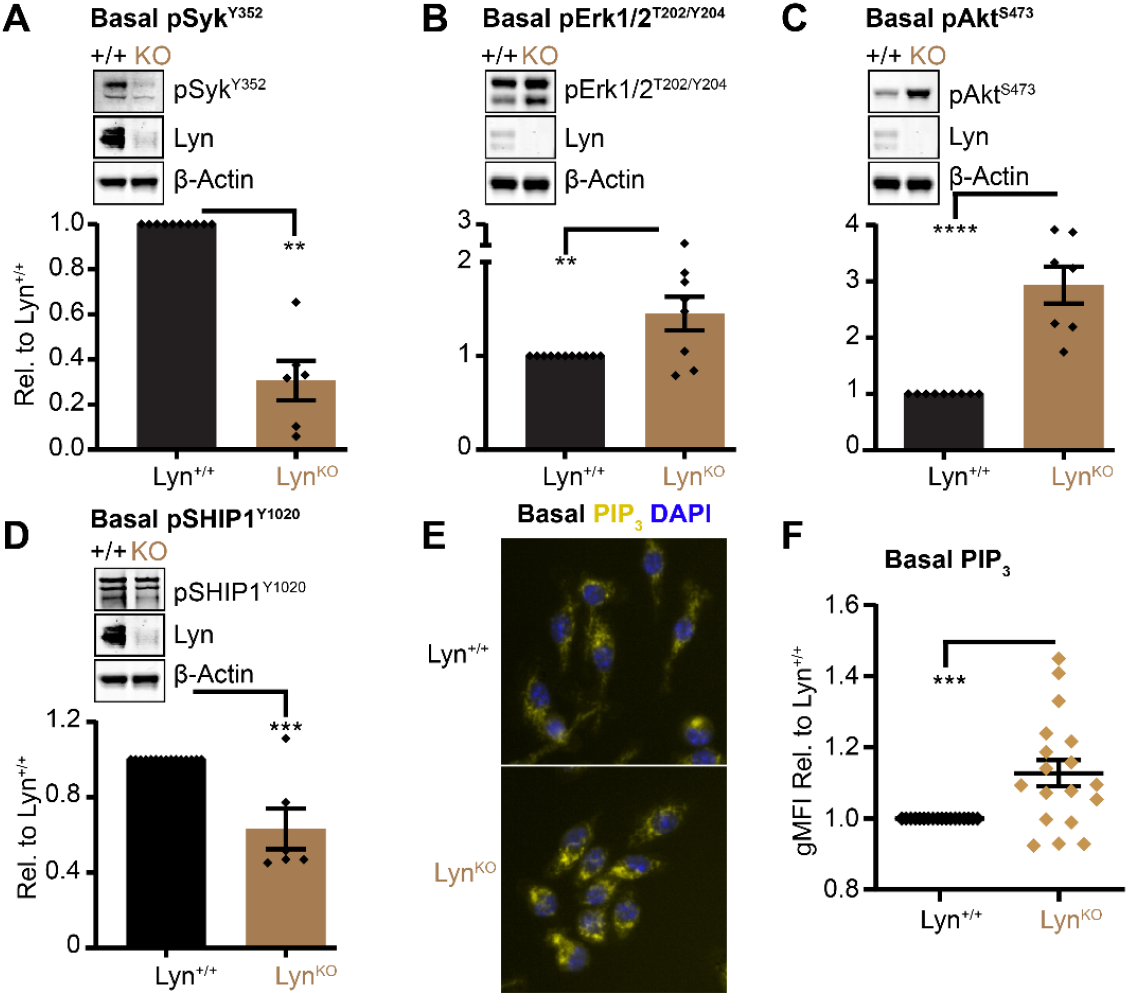
Lyn expression suppresses steady-state phosphorylation of Erk1/2 and Akt and increases phosphorylation of SHIP1 in BMDMs. **(A-D)** Immunoblots and quantifications of lysates from IFN-γ-primed Lyn^+/+^ and Lyn^KO^ BMDMs probed for phosphorylated **(A)** Syk^Y352^, **(B)** Erk1/2^T202/Y204^, **(C)** Akt^S473^, and **(D)** SHIP1^Y1020^. β-Actin is a visual loading control, and quantifications were normalized to total protein content in the gel lane. Data are shown relative to the signal in Lyn^+/+^. n=10-15 for Lyn^+/+^ and 6-7 for Lyn^KO^. Error bars: SEM. Sig. nonparametric, unpaired t with Kolmogorov-Smirnov test, *P=0.0152, **P≥0.007, ***P=0.0006 and ****P≤0.0001. **(E)** Representative images of PIP_3_ (yellow) and nuclear staining (cyan) in IFN-γ -primed BMDMs. **(F)** Quantification of the geometric mean fluorescence intensity (gMFI) of PIP_3_ staining in Lyn^KO^ relative to Lyn^+/+^ BMDMs. Data points reflect the mean and SEM of 3-4 independent wells from 4 biological replicates. Sig. nonparametric, unpaired t test with Kolmogorov-Smirnov test, ^***^P<0.0006.

In myeloid and B cells, Lyn is thought to be the primary SFK that phosphorylates ITIMs and phosphatases, at steady state and after receptor engagement [20, 42]. Given the importance of membrane phospholipids, including PIP_3_ as second messengers for activating the Erk and Akt pathways [43] we posited that a failure to recruit the ITIM-associated inositol phosphatase SHIP1 to the membrane could favor the accumulation of PIP_3_ at steady state and explain the increase in downstream pErk1/2 and pAkt in Lyn^KO^ BMDMs. Lyn^KO^ BMDMs did have a 50% reduction in basal pSHIP1^Y1020^ **(Fig.4D)**, suggesting that SHIP1 recruitment to the membrane and access to its substrate, PIP_3_ [14, 44], is impaired in the absence of Lyn. We confirmed that the reduced SHIP1^Y1020^ phosphorylation in the absence of Lyn was accompanied by an increase in steady-state PIP_3_ **(Fig.4E,F)**.

We have previously shown that splice-form-specific Lyn knockout, LynA^KO^ and LynB^KO^, have distinct effects on myeloproliferation and autoimmunity in mice [23]. We therefore tested whether LynA and LynB contribute differentially to steady-state signal regulation of BMDMs. Expression of LynA or LynB alone at a physiological level (LynA^KO^LynB^+/-^ “LynA^KO^” or vice versa) rescued basal phosphorylation of SHIP1^Y1020^ **(Fig.S1A)**, restoring reduced basal phosphorylation of Erk **(Fig.S1B)** and Akt^S473^ **(Fig.S1C)** to Lyn^+/+^ levels. This suggests that LynA and LynB have overlapping roles in facilitating SHIP1 recruitment to the membrane and phosphorylation, thereby restricting phosphorylation of Erk and Akt. These data demonstrate that Lyn regulates steady state signaling in macrophages by promoting SHIP1 phosphorylation and suppressing accumulation of PIP_3_, which restricts downstream phosphorylation of Akt and Erk. Either splice form of Lyn is sufficient to support SHIP1 signaling and mitigate Akt and Erk hyperphosphorylation.

### Lyn promotes the Erk pathway and restricts the Akt pathway during receptor-independent SFK activation, independently of increased SHIP1 phosphorylation

Having established that Lyn suppresses basal generation of PIP_3_ and activation of the Erk and Akt pathways in macrophages, we interrogated Lyn regulation of signaling following SFK activation in the presence or absence of microscale receptor engagement. The absence of Lyn expression reduced pSyk^Y352^ 3-fold after treatment of Csk^AS^ BMDMs with either the pharmacological SFK activator 3-IB-PP1 **(Fig.5A)** or with particulate depleted zymosan **(Fig.5B)**. Despite the steady-state upregulation of pErk1/2 in Lyn^KO^ cells, induction of pErk1/2 following pharmacological SFK activation was abrogated in the absence of Lyn expression **(Fig.5C)**, consistent with our earlier, qualitative observations [34]. Expression of either LynA or LynB was sufficient to restore Erk activation **(Fig.S1D)**.

**Fig.5:**
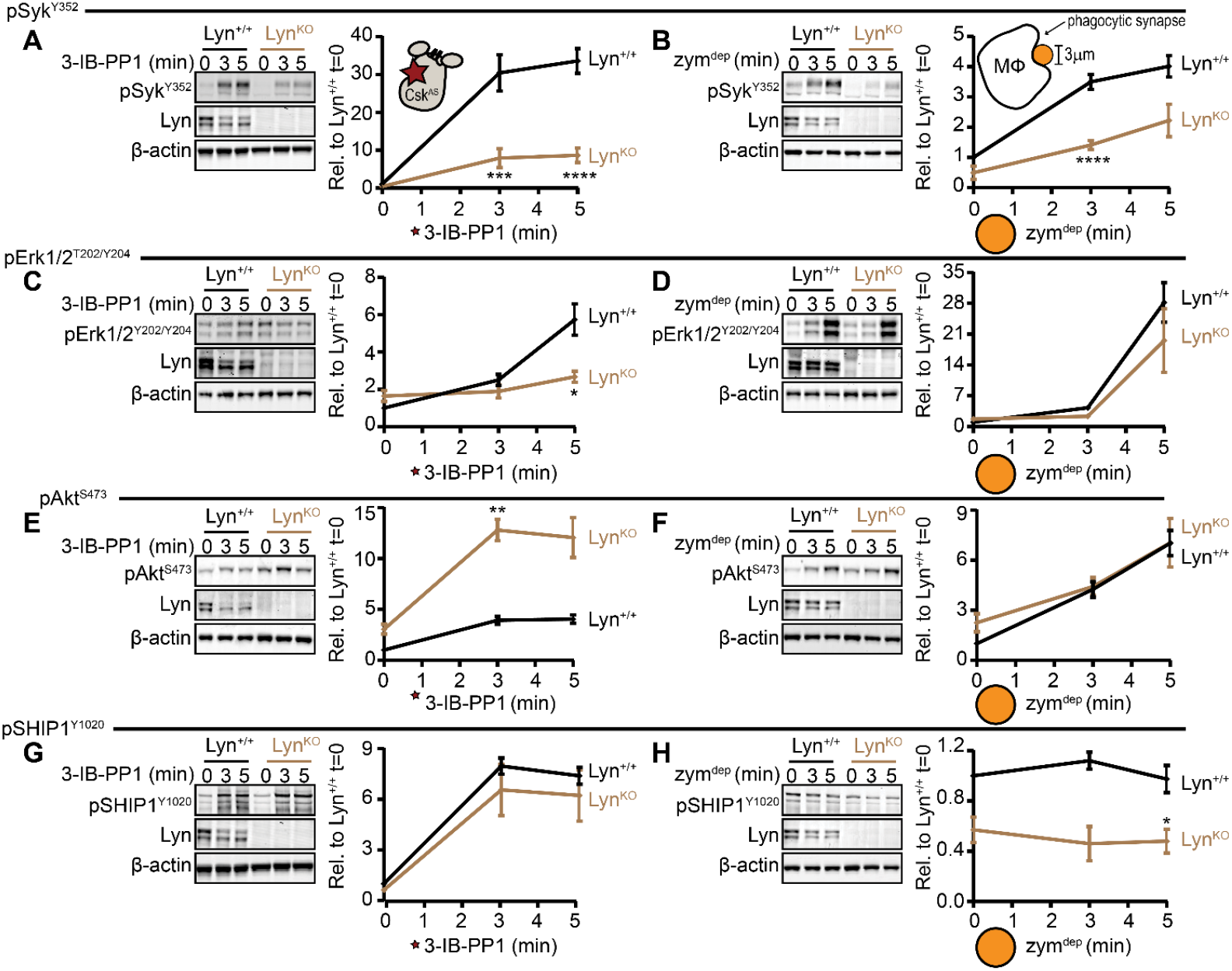
Microscale Dectin-1 engagement relieves a requirement for Lyn to balance activation of the Erk and Akt pathways. Immunoblots and quantifications of **(A-B)** pSyk^Y352^, **(C-D)** pErk1/2^T202/Y204^, **(E-F)** pAkt^S473^, and **(G-H)** pSHIP1^Y1020^ in IFN-γ-primed Csk^AS^ (Lyn^+/+^) or Csk^AS^Lyn^KO^ (Lyn^KO^) BMDMs treated with **(A, C, E, G)** 3-IB-PP1 or **(B, D, F, H)** zym^dep^, corrected for total protein in each gel lane and shown relative to t=0 of Lyn^+/+^; n=10-11 for Lyn^+/+^, 4-7 for Lyn^KO^. Error bars: SEM. Sig. via two-way ANOVA with Sidak’s multiple comparisons test: ^*^P = 0.0143, ^**^P = 0.0041, ^****^P<0.0001.

The absence of Erk phosphorylation in Lyn^KO^ BMDMs following 3-IB-PP1 treatment mirrors the lack of Erk signaling observed in response to the yeast-derived soluble β-glucan Imprime **(Fig.S2A)**. IFN-γ-dependent upregulation of Lyn partially, but not completely, enabled Erk pathway activation via soluble β-glucans. This partial reversal in signaling blockade is not due to contribution from other receptors, as Dectin-1 expression was necessary for signaling **(Fig.S2B)**. Furthermore, binding studies confirmed that the concentration of soluble β-glucans we used saturates at least 80% of Dectin-1 receptors on the macrophage cell surface **(Fig.S2C)**, suggesting that the failure of the Erk response was not due to low Dectin-1 binding. In contrast, the number of depleted zymosan particles per cell in these experiments (10:1) only occupied about 10% of receptors, but it still amplified signaling from a subtler initiating event, further suggesting that maximizing receptor occupancy by nanoparticulate ligands do not lead to productive signaling downstream.

The signaling profile in the presence of soluble β-glucans mirrors pharmacological SFK activation via 3-IB-PP1—in both cases, Lyn expression and inflammatory IFN-γ priming are essential for activation of the Erk pathway [34], but a phagocytic synapse is required for maximal signal amplification of the Erk pathway downstream of SFK activation. This suggests that the particulate “size sensor” function of Dectin-1 does not stem from maximization of receptor occupancy or membrane-proximal signaling, but rather from protecting against the activation of negative regulators, such as SHIP1, and enabling Ras activation and second-messenger pathways to promote downstream Erk signaling. Interestingly, despite the impaired phosphorylation of Syk^Y352^ in Lyn^KO^ BMDMs treated with depleted zymosan, Erk phosphorylation was not impaired **(Fig.5D)**, suggesting Lyn-independent signal amplification downstream of Syk.

In contrast to Erk1/2, we found that the enhanced steady-state phosphorylation of Akt was even further upregulated in 3-IB-PP1-treated Lyn^KO^ BMDMs **(Fig.5E)**. The expression of LynA or LynB was sufficient to restrict Akt^S473^ phosphorylation **(Fig.S1E)**. The formation of a phagocytic synapse and a microscale signalosome, however, balanced activation of the Erk and Akt pathways in a Lyn-independent manner **(Fig.5F)**.

Surprisingly, despite the profound hyperphosphorylation of Akt^S473^ in 3-IB-PP1-treated Lyn^KO^ BMDMs, there was no deficit in phosphorylation of SHIP1^Y1020^ **(Fig.5G)**. This uncoupling of Lyn from SHIP1 phosphorylation upon pan-SFK activation was also surprising, considering the defect in SHIP1^Y1020^ phosphorylation in Lyn^KO^ cells at steady-state. This suggests that the regulation of Akt is dominated by Lyn at steady state, despite the ability of other active SFKs to regulate SHIP1 phosphorylation in the absence of a phagocytic synapse.

The formation of a phagocytic synapse, however, maintained the steady-state differences in pSHIP1^Y1020^ but appeared to protect SHIP1 from further phosphorylation **(Fig.5H)**. In contrast to pan-SFK activation, treatment with depleted zymosan resulted in comparable Akt phosphorylation in the absence of Lyn, reinforcing the premise that the phagocytic synapse protects the signalosome from negative regulation by SHIP1. This protection from SHIP1 could explain how relatively weak activation of Syk could be effectively amplified within the phagocytic synapse to produce normal Erk and Akt pathway activation.

Together, these results suggest that the primary function of Lyn is at steady state and during bursts of SFK activation in the absence of a phagocytic synapse, Lyn supports signal transduction from activated Syk to the Erk pathway (i.e., the Lyn checkpoint [34]) but suppresses excessive activation of Akt and maintains pathway balance.

### Lyn differentially regulates PLCγ2 and PI3K phosphorylation following pharmacological SFK activation

We have shown previously that in the absence of Lyn, the receptor-independent activation of SFKs results in impaired Erk but enhanced Akt phosphorylation. We therefore investigated whether this pathway imbalance stems from differential activation of PLCγ2 and PI3K, which metabolize and phosphorylate PI(4,5)P_2_, respectively, to generate second messengers that amplify signaling downstream. SFK activation in the absence of a phagocytic synapse led to a 2-fold reduction in phosphorylation of PLCγ2^Y1217^ in the absence of Lyn expression **(Fig.6A)**, mirroring the loss of upstream Syk phosphorylation. As with Erk phosphorylation, the expression of LynA or LynB was able to restore PLCγ2^Y1217^ phosphorylation **(Fig.S3)**. PI3K phosphorylation, in contrast, was not affected by loss of Lyn expression **(Fig.6B)**. Interestingly, enhanced Akt phosphorylation in Lyn^KO^ BMDMs suggests that there is PIP_3_ enrichment with 3-IB-PP1 treatment. However, even though Lyn^KO^ BMDMs had elevated levels of PIP_3_ at steady state, PIP_3_ levels were depleted with 3-IB-PP1 treatment **(Fig.6C)**. Interestingly, Lyn^KO^ BMDMs still had marginally higher PIP_3_ levels with 3-IB-PP1 treatment. This lack of PIP_3_ enrichment is potentially a result of strong activation of SHIP1 generated in both genotypes by receptor-independent SFK activation.

**Fig.6:**
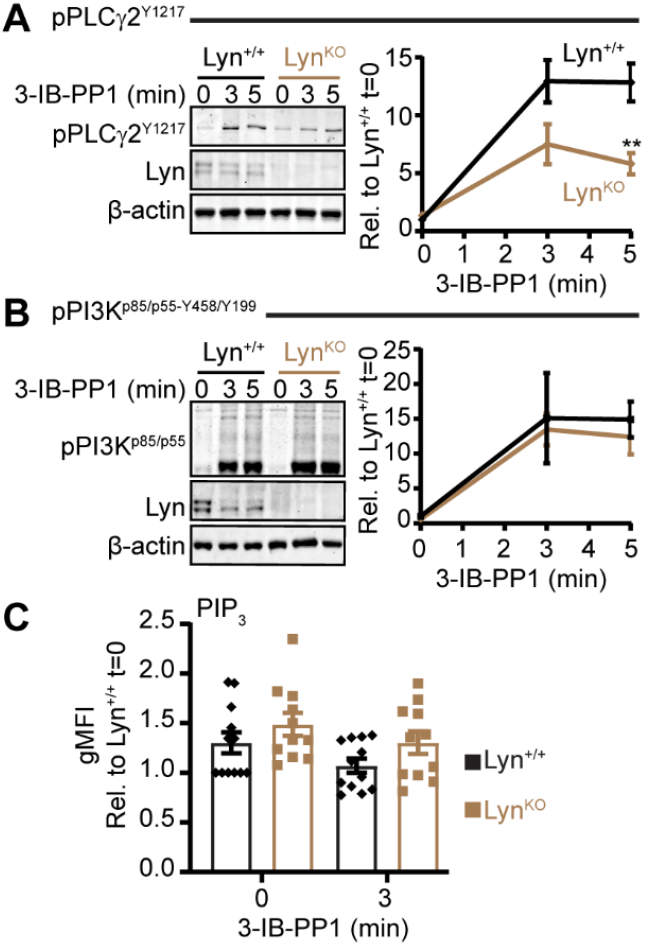
Lyn is required for maximal phosphorylation of PLCγ2 but not PI3K. Immunoblots and quantification of phos-phorylated **(A)** PLCγ2^Y1217^ and **(B)** PI3K^p85/p55-Y458/Y199^ in Lyn^+/+^ and Lyn^KO^ BMDMs treated with 3-IB-PP1, corrected for the total protein content in each lane and shown relative to t=0 of Lyn^+/+^. In all panels error bars: SEM. A-B show n=10 for Lyn^+/+^ and 3-4 for Lyn^KO^. **(C)** Immunofluorescence-microscopy-derived geometric mean fluorescent intensity (gMFI) measured by immunofluorescence, of total cellular PIP_3_ via immunofluorescence, shown relative to Lyn^+/+^ t=0; n=9-12 replicates from cells prepared from 3 mice. Sig.: two-way ANOVA with Sidak’s multiple comparisons test: ^**^P=0.0015.

Here we demonstrate that while Lyn is essential for optimal PLCγ2 phosphorylation, other kinases can support PI3K activation. This Lyn-independent PI3K activity, combined with already elevated steady-state PIP_3_ and pAkt^S473^, may help maintain Akt pathway function but the lack of PIP_3_ enrichment does not fully account for Akt hyperactivation.

### The SFKs Hck and Fgr do not assist Lyn in maintaining steady-state phosphorylation of SHIP1 or restricting Akt hyperphosphorylation during pharmacological SFK activation

We showed that Lyn regulates steady-state signaling by phosphorylating SHIP1 and restricting Akt phosphorylation. Since Csk^AS^ inhibition induces activation of all the SFKs, including Hck and Fgr [34], we investigated the contributions of Hck and Fgr to signaling at steady state and following pharmacological SFK activation. In contrast to Lyn^KO^ BMDMs, Hck and Fgr double-knockout (Hck^KO^Fgr^KO^) BMDMs **(Fig.S4A,B)** had no defect in steady-state phosphorylation of SHIP1^Y1020^ **(Fig.7A)**. Although the pSHIP1^Y1020^ levels were comparable between Hck/Fgr-expressing and Hck^KO^Fgr^KO^ BMDMs, we did observe a slight increase in basal pAkt^S473^ in Hck^KO^Fgr^KO^ BMDMs **(Fig.7B)**. However, this increase was not at the same magnitude as Lyn^KO^ BMDMs. In contrast to our findings with Lyn^KO^ BMDMs, there was no significant elevation of pAkt^S473^ seen in Hck^KO^Fgr^KO^ BMDMs with the addition of 3-IB-PP1.

**Fig.7:**
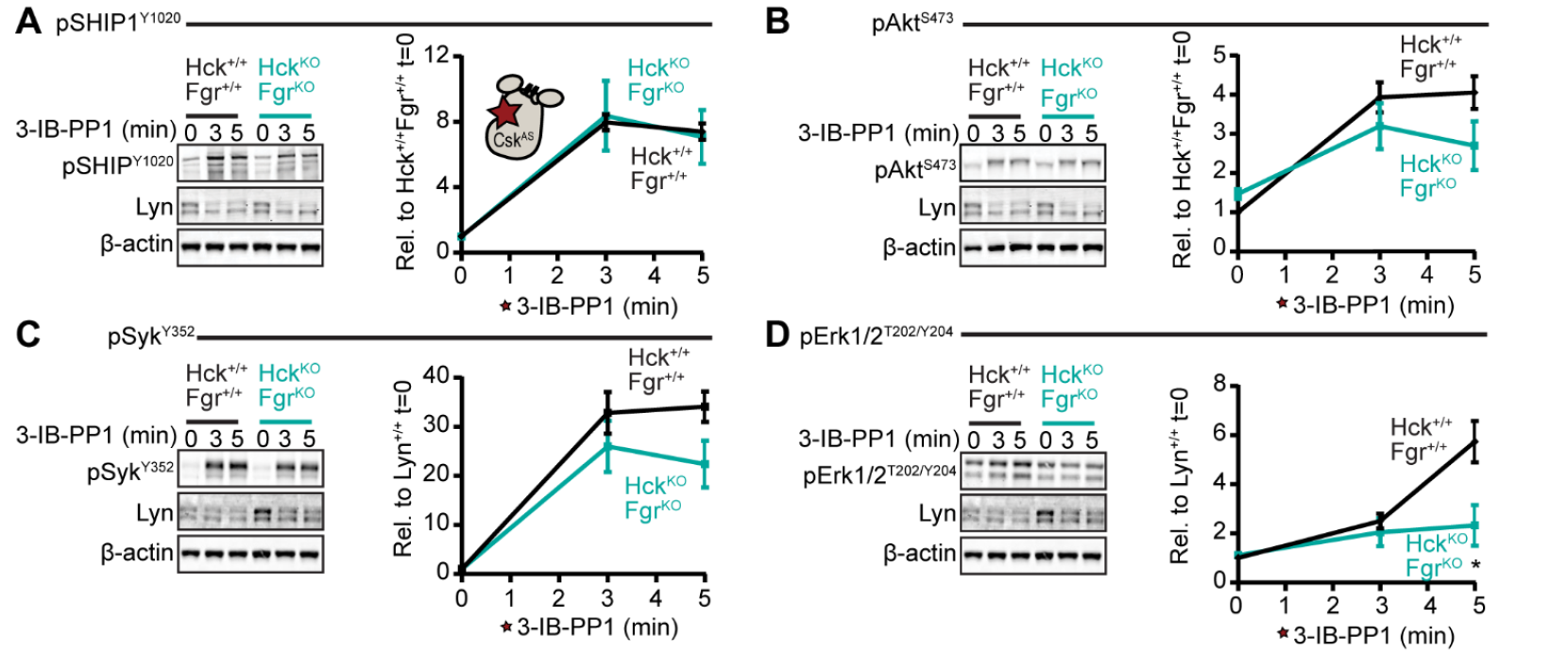
Lyn functions uniquely in regulating the Akt pathway but not the Erk pathway in steady-state and SFK-activated macrophages. Immunoblots and quantification of **(A)** pSHIP1^Y1020^, **(B)** pAkt^S473^, **(C)** pSyk^Y352^, and **(D)** pErk^T202/Y204^ in CskAS (Hck^+/+^Fgr^+/+^) and Csk^AS^Hck^KO^Fgr^KO^ (Hck^KO^Fgr^KO^) BMDMs, corrected for the total protein in each lane and shown relative to t=0 of Hck^+/+^Fgr^+/+^. Error bars: SEM, n=10-11 for Hck^+/+^Fgr^+/+^ and n=4 for Hck^KO^Fgr^KO^. Sig. 2-way ANOVA with Sidak’s multiple comparisons test: ^*^P = 0.0499.

### Expression of Lyn, Hck, and Fgr are necessary for Erk phosphorylation following 3-IB-PP1 treatment

In contrast to complete Lyn^KO^ BMDMs, we observed that Syk^Y352^ phosphorylation at steady state was not impaired by the lack of Hck and Fgr **(Fig.7C)**. Similarly, there was no decrease in Syk phosphorylation after pan-SFK activation in the absence of a phagocytic synapse. This showed that Lyn kinase plays a predominant role in signal initiation downstream of activated SFKs in macrophages.

Steady-state Erk phosphorylation in Hck^KO^Fgr^KO^ BMDMs was comparable to Hck/Fgr-expressing cells **(Fig.7D)** and Syk phosphorylation was intact in Hck^KO^Fgr^KO^ BMDMs after pan-SFK activation. Despite these consistencies with Hck/Fgr-expressing cells, Erk phosphorylation was impaired in Hck^KO^Fgr^KO^ BMDMs, comparable to Lyn^KO^ BMDM levels. This suggests that in signaling nanoclusters Lyn plays a crucial role in initiating and transducing signals from Syk to PLCγ2. However, the overall strength of activated SFKs ultimately determines Erk induction, independently of upstream signaling.

## Discussion

Signaling outcomes in myeloid cells and lymphocytes are determined by the valency of ITAM receptor ligands and the surface area of receptor engagement. Low-valency (nanoscale) ligation of Dectin-1 by soluble β-glucans does not pass the required threshold to initiate signaling, phagocytosis and release of reactive oxygen in macrophages [31]. Dectin-1 also binds to annexin core domains in apoptotic cells via an orthogonal binding site [45]. This lower-valency Dectin-1 clustering by apoptotic cells leads to some activation of Syk but failure to activate downstream factors such as NF-κB, thereby promoting immune tolerance. In contrast, canonical, microscale engagement of Dectin-1 by depleted zymosan particles drives NF-κB activation and pro-inflammatory signaling [45]. Similarly, in B cells, soluble antigens with low-valency receptor engagement, induce membrane-proximal signaling but fail to propagate signals downstream to Erk and Akt, in contrast to highly multivalent particulate antigens. This lack of signal amplification is likely due to the activation of ITIM-associated phosphatases and limited access of PLCγ2 to PI(4,5)P2, which together impair second-messenger generation and down-stream signal amplification [46].

Here we show that pharmacological, receptor-independent activation of SFKs in macrophages results in dispersed, nanoscale pSyk^Y352^ signaling clusters without the semi-continuous, microscale signalosome characteristic of a phagocytic synapse. These signaling nanoclusters generate robust membrane-proximal signaling but largely fail to propagate to second-messenger-dependent downstream signaling, even after inflammatory priming with IFN-γ. This mirrors the defect in downstream signal amplification observed in B cells exposed to soluble antigens [46] and 3-IB-PP1-treated thymocytes and B cells [36, 43]. This failure of signal amplification may be attributed to enhanced phosphorylation of the ITIM-associated phosphatases SHIP1 and SHP-1, which restrict signaling in the absence of a phagocytic synapse. However, despite the elevated phosphorylation of the tyrosine phosphatase SHP-1 following 3-IB-PP1 treatment, we did not observe bulk dephosphorylation of tyrosine-phosphorylated proteins such as Syk, PLCγ2, and PI3K. Interestingly, enhanced SHIP1 phosphorylation was associated with rapid PIP_3_ depletion, similar to what has been observed in B cells subjected to pharmacological SFK activation [43]. This suggests that B cells and macrophages share a common regulatory mechanism to limit Erk and Akt activation in response to abrupt SFK disinhibition or engagement of soluble receptor ligands, highlighting the crucial role of SHIP1 in distinguishing pathogenic particle encounters from inappropriate activation of macrophages.

Previous studies have shown that steady-state SFK and ITAM activity can either potentiate or inhibit myeloid-cell responses downstream of ITAMs, Toll-like receptors, and cytokine receptors [16, 38, 39]. Here we show that IFN-γ-dependent up-regulation of Lyn can sensitize macrophages to signaling in response to low-valency, soluble β-glucans. Our findings extend these observations to reveal a unique role for Lyn in steady-state regulation. Lyn activity is essential for maintaining tonic phosphorylation of SHIP1, limiting the accumulation of PIP_3_, and suppressing basal Erk and Akt signaling. Intriguingly, the net function of Lyn in Erk activation during pharmacological SFK activation or signaling in response to soluble β-glucans is positive-regulatory [34]. In contrast to Erk activation and previous observations in B cells [43], pharmacological activation of SFKs in Lyn^KO^ BMDMs led to enhanced Akt phosphorylation. This regulation of Akt phosphorylation in the absence of a phagocytic synapse is unique to Lyn as Hck and Fgr double knockout BMDMs had no changes in basal or induced Akt phosphorylation with 3-IB-PP1. Interestingly, we did not observe hyperphosphorylation of PI3K [47] or impaired SHIP1 phosphorylation in Lyn^KO^ BMDMs following pan-SFK activation that explains the heightened Akt response. This observation reinforces Lyn’s dominant role in restricting Akt hyperresponsiveness at steady-state, preventing spurious Akt activation and maintaining pathway balance during disinhibited SFK activation, in the absence of a phagocytic synapse.

In contrast to 3-IB-PP1-treated BMDMs, SHIP1 and SHP-1 are not phosphorylated above steady-state levels after microscale engagement of Dectin-1, a profoundly different outcome than the levels of activating phosphorylation on Syk. Time-dependent phosphorylation of SHIP1 further confirms that during the formation of a phagocytic synapse, SHIP1 is excluded from the signalosome and thus protected from SFK-dependent phosphorylation, preventing negative feedback. Furthermore, particulate β-glucans override the negative regulatory role of Lyn and restore the balance of Erk and Akt signaling.

Uncoupling SFK activation from the formation of a phagocytic synapse has enabled us to assess how macrophages resist fluctuations in SFK activity, which can occur stochastically or due to low-valency receptor interactions with soluble ligands, while simultaneously being ready to mount strong, potentially tissue-toxic responses to genuine encounters with microscale pathogens. While the exclusion of the membrane phosphatases CD45 and CD148 are implicated in tuning the sensitivity of macrophages to soluble or particulate Dectin-1 ligands [1, 31], our work has shown that tyrosine dephosphorylation of membrane-proximal signaling proteins is not necessarily the key point of signaling blockade. Rather, our findings suggest that exclusion of the cytosolic phosphatases SHIP1 and SHP-1 from the phagocytic synapse allows upstream tyrosine-phosphorylation and activation of membrane-proximal signaling to connect productively with second-messenger generation and downstream Ras, Erk, and Akt. These observations are consistent with a model in which Lyn and SHIP1 serve as gatekeepers of macrophage activation, maintaining signal homeostasis and preventing inappropriate activation in response to low-valency or soluble stimuli. Thus, the formation of a phagocytic synapse allows bypass of these negative-regulatory functions and produce amplified and balanced Erk and Akt activation.

## Materials and Methods

### Mice

All mice were derived from a C57BL/6 background. Csk^AS^Lyn^KO^ and Csk^AS^Hck^KO^Fgr^KO^ mice were generated by crossing Csk^KO^Csk^AS^-transgenic mice with the appropriate Csk^+/-^ knockout [23, 34]. LynA^KO^ and LynB^KO^ mice were used as F1 crosses of Lyn^KO^ and Lyn^[A or B]KO^(CRISPR) mice to ensure WT-like expression of the remaining Lyn isoform, as reported previously [23]. Animal studies were performed in compliance with the policies of the University of Minnesota, the American Association for Accreditation of Laboratory Animal Care, and National Institutes of Health, under Animal Welfare Assurance no. A3456-01 and Institutional Animal Care and Use Committee protocol no. 2209-40372A. All animals were housed in specific-pathogen-free conditions and provided with care by licensed veterinarians.

### Preparation of BMDMs

BMDMs were prepared as previously described [34]. Bone-marrow was flushed from the tibia and femurs of mice and subjected to hypotonic erythrocyte lysis. The bone marrow was then plated in non-tissue-culture-treated plates (Corning, VA) in Dulbecco’s Modified Eagle Medium (Corning Cellgro, Manassas, VA, USA) supplemented with 10% heat inactivated FBS (Omega Scientific, Tarzana, CA), 0.11 mg/ml sodium pyruvate (Corning, VA), 2 mM penicillin/streptomycin/L-glutamine (Sigma-Aldrich, St. Louis, MO, USA), and 10% CMG-14-12-cell–conditioned medium as a source of macrophage colony-stimulating factor (M-CSF). After 7 days of culture, BMDMs were lifted off the plate using 5 mM EDTA and re-plated at 1M cells/well in non-treated 6-well plates (Corning, VA, USA) in unconditioned medium with 25 U/ml IFN-γ (Peprotech, Rocky Hill, NJ, USA).

### RNA extraction

Blood from either WT or Hck Fgr double-knockout mice were collected into tubes with EDTA, and RNA was extracted according to the directions in RiboPure-Blood kit, Cat# AM1928 (Life technologies, Carlsbad, CA, USA). cDNA synthesis done using qScript cDNA synthesis kit, Cat# 95047 (Quantabio, Beverly, MA, USA). SYBR green was used to detect Fgr in 1 µg total RNA (forward: CTCAAGGCCGGACTTCGT, reverse: TGCTTCTCATGTTGCCAG-TGTT) and Cyclophilin amplification (forward: TGCAGGCAAAGACACCAATG, reverse: GTGCTCTCCAC-CTTCCGT).

### Cell stimulation

BMDMs were treated with 10 µM 3-IB-PP1, 10 particles/cell depleted zymosan (Sigma-Aldrich), or 0.8 µg/ml of the yeast-derived soluble β-glucan Imprime PGG [48] (Patent #: US11815435B2) (Biothera, Inc., Eagan, MN, USA) via pulse spinning and placed at 37°C until the timepoints were completed. Signal propagation was suspended by placing the plate on ice. Cells were lysed in sodium dodecyl sulfate (SDS) buffer (128mM Tris base, 10% glycerol, 4% SDS, 50mM dithiothreitol (DTT), pH 6.8). The cells were scraped using cell scrapers and incubated at 37°C for 5 minutes. The lysates were collected into 1.5 ml microcentrifuge tubes and subjected to sonication via Diagenode Bioruptor (Diagenode Inc, Denville, NJ, USA) at 50% duty for 3 minutes to shear DNA. The lysates were then boiled for 15 min and stored at -20°C.

### Quantification of Ras-GTP

After treatment with 3-IB-PP1 or depleted zymosan, wells were washed with ice-cold 1x phosphate-buffered saline (PBS) and lysed in buffer provided by the Active Ras Detection kit, Cat# 8821s (Cell signaling Technology, Danvers, MA, USA) supplemented with HALT phosphatase and protease inhibitors, Cat# 78440 (Thermo Fisher Scientific, Waltham, MA, USA). Equivalent amounts of protein were used for subsequent steps, per manufacturer instructions.

### Immunoblotting

Samples were separated on a 7% NuPage Tris-Acetate gel (Invitrogen, Carlsbad, CA, USA) and then transferred to Immobilon-FL PVDF membrane (EMD Millipore, Burlington, MA, USA). Revert Total Protein stain (TPS, LI-COR Biosciences, Lincoln, NE, USA) was used to quantify the total protein content in each lane. After TPS removal, the membrane was cut as appropriate and blocked 1 h with TBS Intercept Blocking Buffer (LI-COR, Lincoln, NE, USA). The membranes were then incubated overnight at 4°C with the appropriate primary antibodies purchased from Cell Signaling Technologies. The membranes were visualized using an Odyssey CLx near-infrared imager (LI-COR, Lincoln, NE, USA) after 1.5-hour incubation with secondary antibodies purchased from LI-COR Biosciences. Densitometry quantification was performed in ImageStudio (LI-COR), as described previously [49]. Antibodies were obtained from Cell Signaling Technologies (rabbit anti-pSyk^Y352^ 65E4 ID/catalog # (#)2717, rabbit anti-pPLCγ^Y1217^ #3871, rabbit anti-pSHIP1^Y1020^ #3941, rabbit anti-pSHP-1^Y564^ D11G5 #8849, rabbit anti-pErk1/2^T202/Y204^ D13.14.4e #4370, rabbit anti-pAkt^S473^ 193H12 #4058, rabbit anti-pPI3K^p85(Y458)/p55(Y199)^ #4228, mouse anti-β-Actin 8H10D10 #3700), Abcam (Cambridge, United Kingdom) (mouse anti-LynA+B Lyn01 ab1890), Santa Cruz (Dallas, TX, USA) (rabbit anti-LynA+B 44 #sc-15, goat anti-Hck M-28 #sc-1428), and LICOR (donkey anti-mouse IgG 800CW #925-32212, donkey anti-rabbit IgG 800CW #925-32213, donkey anti-mouse IgG 680LT #925-68022, donkey anti-goat IgG 680LT #926-32214).

### Dectin-1 binding competition assay

After BMDM differentiation, cells were plated in nontreated 6 well plates in unconditioned media with or without 25 U/ml IFN-γ. Cells were treated with varying concentrations of Imprime PGG or depleted zymosan to saturate Dectin-1 receptors. Following β-glucan treatment, cells were incubated with a 1:10 dilution of anti-mouse Dectin-1-APC antibody (Cat# 130-102-250, Miltenyi Biotec, Gaithersburg, MD, USA). Binding was assessed by measuring the decrease in gMFI values compared to untreated BMDMs. gMFI values were corrected for background fluorescence and plotted against the corresponding β-glucan concentration using GraphPad Prism (San Diego, CA). Data points were fitted using a nonlinear one-phase decay model. The concentration of soluble or particulate β-glucans required to saturate Dectin-1 receptors was modeled using the equation: Y = (Y_0_ - Plateau) * e^(−KX) + Plateau.

### Confocal microscopy and image analysis

BMDMs were plated onto 15 mm glass coverslips, Cat# CLS-1763-015 (Chemglass Life Sciences, Vineland, NJ, USA) overnight in unconditioned medium with 25 U/ml IFN-γ (Peprotech). Cells were treated as indicated at 37°C followed by fixation in 4% PFA for 15 min at RT and washed with 10 mM Tris/PBS buffer. Samples were washed with PBS and then simultaneously blocked and permeabilized with 0.1% Triton X-100/2% BSA/PBS for 10 min at RT. Samples were labeled with rabbit Anti-pSykY352, Cat# ab300398 (Abcam) and mouse anti-Lyn, Cat# ab1890 (Abcam) diluted in 2% BSA/PBS for 60 min at RT, washed three times with PBS, followed by secondary F(ab’)_2_ labeling with anti-Rabbit-AF647, Cat# 111-606-047 (Jackson ImmunoResearch, West Grove, PA), to label pSykY352, and anti-Mouse-AF488, Cat# 115-546-071 (Jackson ImmunoResearch), to label Lyn, for 60 min at RT before DAPI staining and mounting in ProLong Gold, Cat# 9071S (Cell Signaling Technology). Confocal images were obtained using a Leica STELLARIS 8 (Nussloch, Germany) inverted laser scanning confocal microscope using galvonomic scanners and spectral internal HyD-S detectors and a fixed 405nm violet laser (DAPI excitation) and pulsed white-light laser for excitation of AF488 and AF647. A Leica Harmonic Compound PL apochromatic CS2 63X oil objective with a correction collar (1.4 NA) was used for imaging. Laser powers were 5.5% for 488 nm and 6% for 647 nm. Two-hybrid detectors collected photons at 505–565 nm gain 7% (AF488) and 655-723 nm gain 3.5% (AF647). Imaris 9.9.1 software was used to define protein clusters using the Surfaces toolbox of pSyk clusters and of the depleted zymosan particles. The surface area of pSyk^Y352^ clusters and depleted zymosan was calculated using Imaris image analysis software (South Windsor, CT, USA) and plotted using GraphPad Prism.

### Measurement of phosphatidylinositol abundance

Following activation, BMDMs were incubated in 4% PFA at 4°C for 1 h. Cells were washed with PBS with 0.1% Tween 20 (PBST) and permeabilized with 0.1% Triton X-100 for 10 min. Cells were then blocked 1 h in 10% FBS before staining with an antibody against PIP_3,_ Cat# Z-B3345B (Echelon Biosciences, Salt Lake City, UT) in PBST for 1 h at RT. Cells were washed with PBST and incubated 1 h with a secondary streptavidin AF547, Cat# S32356 (Thermo Fisher Scientific). Cells were also visualized by staining with DAPI, Cat# D9542 (Sigma-Aldrich, St. Louis, MO) and phalloidin, Cat# A12379 (Thermo Fisher Scientific). Finally, cells were washed with PBST and analysis of the fluorescence intensity was performed on a Thermo Fisher CX5 imager. Object detection was performed using the phalloidin stain and a mean fluorescent intensity for PIP_3_ was determined for each well.

## Acknowledgements

Many thanks to Dr. Kevan Shokat (UCSF) for supplying us with the Csk^AS^ inhibitor 3-IB-PP1, to Dr. Clifford Lowell (UCSF) for the gift of Hck^KO^Fgr^KO^ mice, and to Biothera, Inc. (Eagan, MN), for donating Imprime PGG β-glucans for this study.

## Data availability

The data that support the findings of this study are available from the corresponding author upon reasonable request.

## Author Contributor Roles

Conceptualization-Lead (SES, TSF), Data Curation-Lead (SES, TSF), Formal analysis-Lead (SES, TSF), Formal analysis-Supporting (WKK, RTC, WFH), Funding acquisition-Lead (TSF), Funding acquisition-Supporting (WKK, WFH), Investigation-Lead (SES, TSF), Investigation-Supporting (WKK, RTC, WLS, STR, MGN), Methodology-Lead (SES, TSF), Methodology-Supporting (WKK, RTC, WFH), Project administration-Lead (TSF), Resources-Lead (TSF), Supervision-Lead (TSF), Supervision-Supporting (SES, WFH), Validation-Lead (TSF), Visualization-Lead (SES, TSF), Visualization-Supporting (WKK), Writing-original draft-Lead (SES, TSF), Writing-original draft-Supporting (WKK, RTC), Writing-reviewing and editing-Lead (SES, TSF), Writing-reviewing and editing-Supporting (WKK, RTC, WLS, STR, MGN, WFH)

## Topic

Molecular and Structural Immunology

## Supplementary Figures

**Fig.S1:**
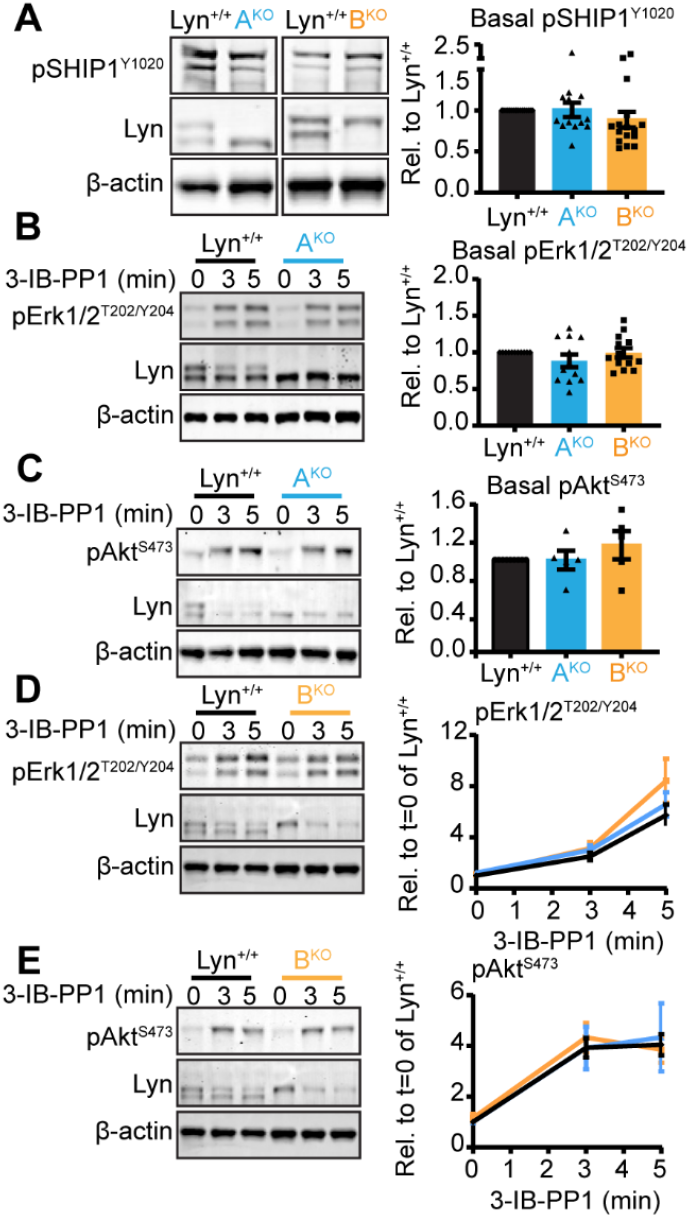
pSHIPl^Y1020^, pErkl/2^T202^/^Y204^ and pAkf^S473^ in LynA^K0^ and LynB^K0^ BMDMs. Immunoblots and quantifications of basally and 3-IB-PP 1-induced phosphoproteins in lEN γ -primcd LynA^K0^ (A^KO^) and LynB^KO^ (B^k0^) BMDMs. Quantifications corrected for total protein and shown relative to Lyn+/+. (A) basal pSHIPl^YI020^, (B) basal pErkl/2^T202^/^Y204^, (C) basal pAkt^S473^, (D) 3-IB-PPI-induccd pErkl/2^T202/Y204^, and (E) 3-IB-PPl-induced pAkt^S473^. Error: SEM, n=9,15 for Lyn+/+, n=4 or14 for LynA^K0^ and n=5 or 15 biological replicates for LynB^KO^. Sig. for basal quantifications: one-way ANOVA with Tukey’s multiple comparisons test. Sig. for induced com-parisons: two-way ANOVA with Tukey’s multiple comparisons test. No significant differences.

**Fig.S2:**
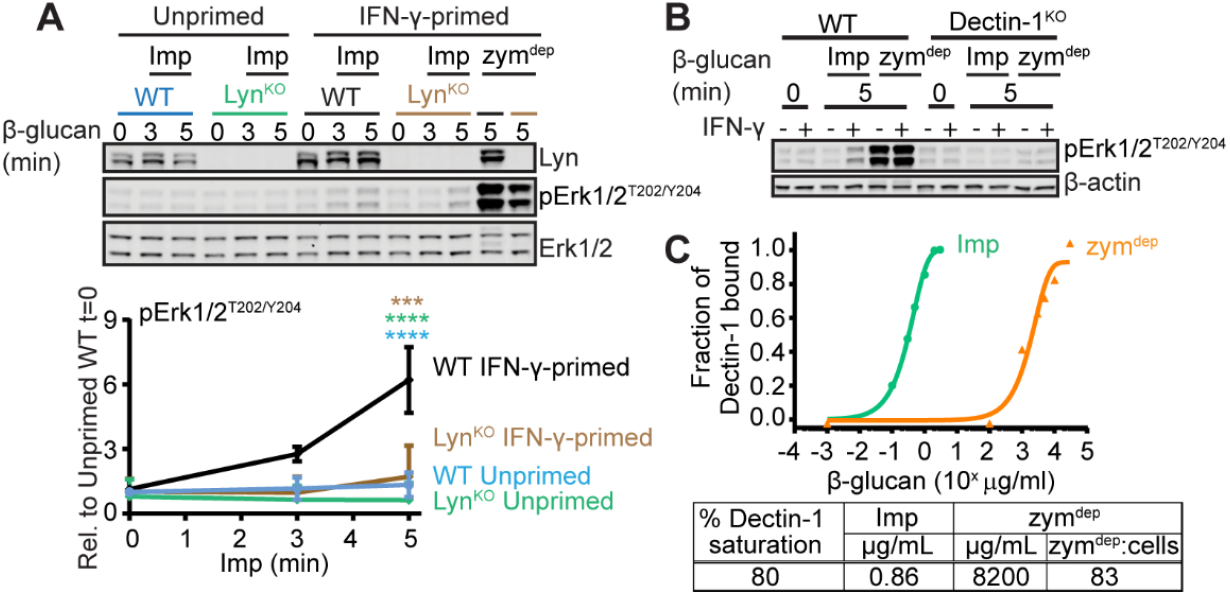
Erk phosphorylation in BMDMs treated with soluble or particulate β-glucans. **(A)** Immunoblols and quanti fiactions of pFrkl/2,T20/Y204in unprimcd or ll-N-γ -primcd WTand Lyn^KO^ BMDMs treated with the soluble β -glucan Imprime PGG(lmp)or particulate 0-glucan zym‘^k^’’, corrected for total protein and shown relative to unprimcd WT t=0. Error: SEM, n= 8 for WT. n=4 for Lyn^KO^. Sig. 2-way .ANOVA with Tukey’s test: ****I*<0.0001 and ***P - 0.0002. (B) Immtinoblot probed for pErkl/ZT202/Y204in WTand Declin-1^kO^ BMDMs treated with eitlter Imp or zym^dep^. (C) Binding competition assessing the treatment dose of Imp or zym^dep^ necessary to produce comparable levels of Dectin-1 saturation on BMD.Ms.

**Fig.S3:**
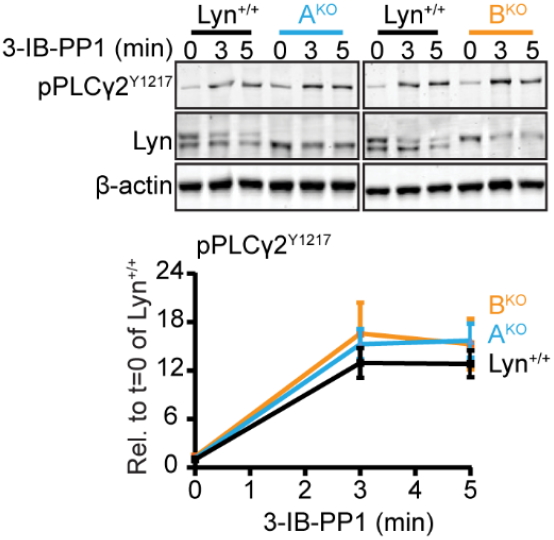
PLCγ2 phosphorylation in LynA^KO^and LynB^KO^ BMDMs with 3-IB-PP1 treatment. Immunoblots and quantifications of pPLCy2^YI217^ in IFN-γ-primcd LynA^K0^ (A^KO^) and LynB^K0^ (B^KO^) BMDMs, shown corrected for total protein stain and relative to t=0 of Lyn^+/+^. Error: SEM, n= 17 for Lyn^+/+^, n=6 for A^KO^, and n=l 1 for LynB^KO^. Sig. two-way ANOVA with Tukey’s multiple comparisons lest. No significant differences found.

**Fig.S4:**
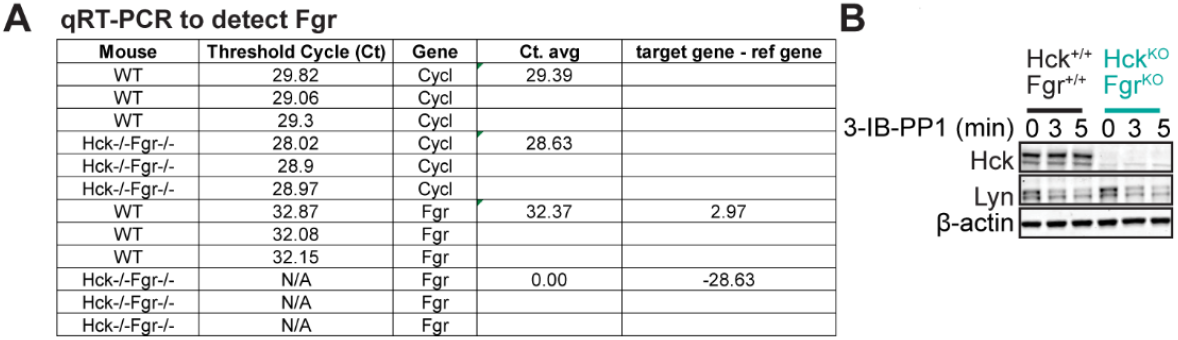
qRT-PCR and western blots for Fgr and Hck: (A) qRT-PCR to detect Fgr in WT and double knockout peripheral blood mononuclear cells. **(B)** Immunoblols probed for lick ard Lyn in WT (Hck^+/+^ Fgr^+/+^) and double knockout BMDMs.

